# SARS-CoV-2 3CLpro mutations confer resistance to Paxlovid (nirmatrelvir/ritonavir) in a VSV-based, non-gain-of-function system

**DOI:** 10.1101/2022.07.02.495455

**Authors:** Emmanuel Heilmann, Francesco Costacurta, Andre Volland, Dorothee von Laer

## Abstract

Protease inhibitors are among the most powerful antiviral drugs. A first protease inhibitor against the SARS-CoV-2 protease 3CLpro, Paxlovid (nirmatrelvir / ritonavir), has recently been authorized by the U.S. FDA for emergency use (EUA 105 Pfizer Paxlovid). To find resistant mutants against the protease-inhibitor-component of Paxlovid, nirmatrelvir, we engineered a chimeric Vesicular Stomatitis Virus (VSV). By replacing an intergenic region, which is essential for separate gene transcription, with 3CLpro, this chimeric VSV became dependent on the protease to process two of its genes. We then applied selective pressure with nirmatrelvir to induce mutations. The effect of those mutants was confirmed by re-introduction in the 3CLpro and testing with a recently developed cellular assay. Furthermore, we found that mutations predicted by our method already exist in SARS-CoV-2 sequence depositions in NCBI and GISAID data bases. These may represent emerging resistant virus variants or a natural heterogeneity in the susceptibility to nirmatrelvir.

**One-Sentence Summary:** Mutations of the main protease of SARS-CoV-2 result in resistance against licensed drugs such as Paxlovid (nirmatrelvir / ritonavir).

**Graphical abstract:** 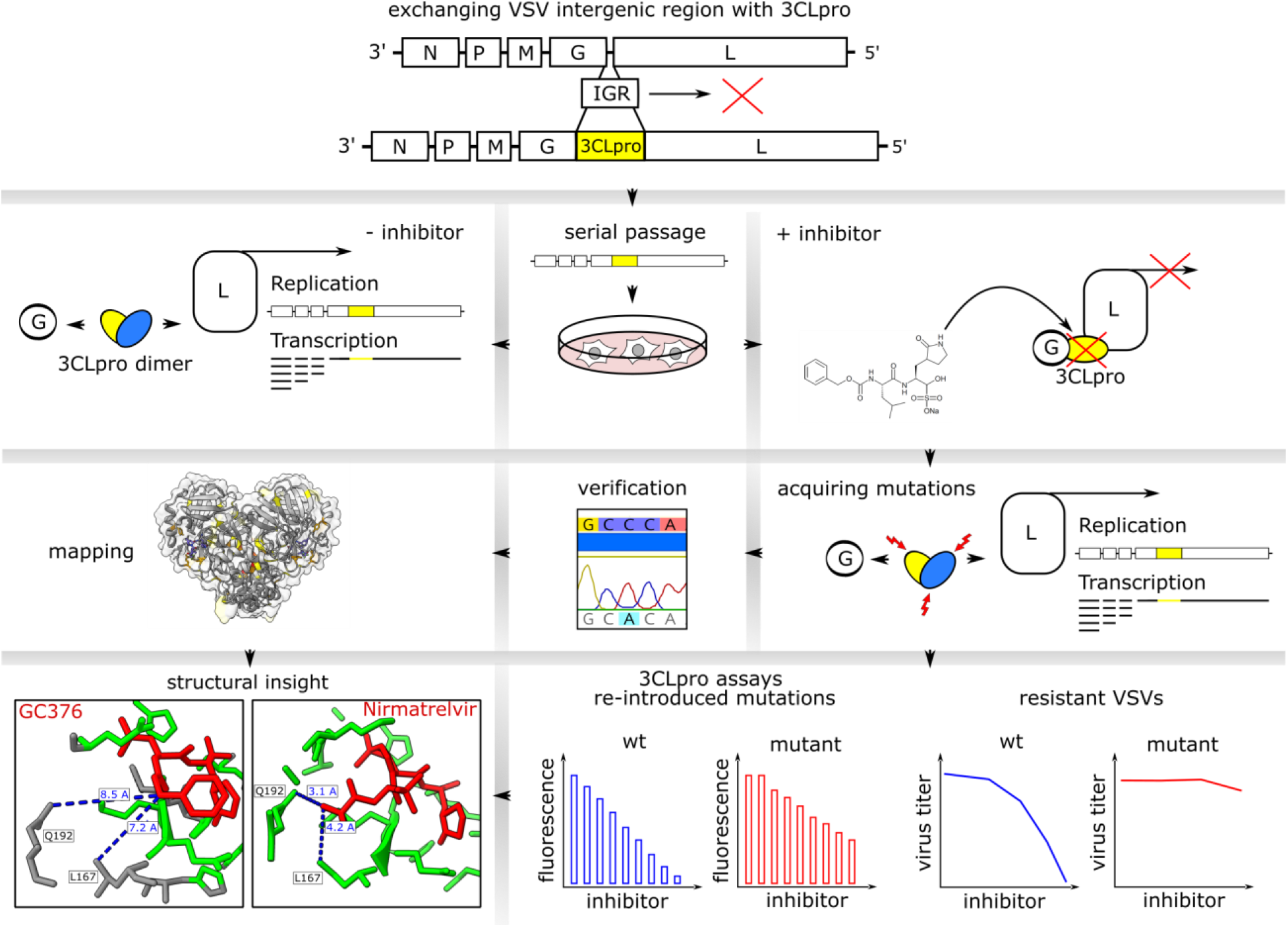

## Main Text

The first protease inhibitor against the SARS-CoV-2 main protease (Mpro, 3CLpro, Nsp5), Paxlovid (nirmatrelvir / ritonavir), has recently been authorized by the U.S. FDA for emergency use in high-risk SARS-CoV-2-infected individuals (EUA 105 Pfizer Paxlovid, 22.12.2021). In the studies leading to the Paxlovid EUA, mouse hepatitis virus (MHV) 3CLpro was used as a surrogate for SARS-CoV-2 3CLpro to generate resistance data. However, for SARS-CoV-2, no resistance data have been published so far. We compared a panel of different viral proteases, containing MHV 3CLpro and SARS-CoV-2 3CLpro in a recently developed gain-of-signal assay (*1*). We found that although nirmatrelvir has broad activity against all tested proteases, MHV 3CLpro showed a significantly weaker response compared to SARS-CoV-2 3CLpro (**Figure 1a**). Indeed, the sequence identity of the two proteins is only 50 % (**Figure S1**). The structures of the SARS-CoV-2 and MHV 3CLpro enzymes are strongly conserved. However, the interaction site of nirmatrelvir (a distance of 5 ångström or less from the compound) shows seven amino acid differences between the two enzymes, namely H164-Q, M165 - L, P168 - S, V186 - R, R188 - A, T190 - V and A191 - V (counting from the first residue (serine) after the glutamine of the N-terminal cleavage site). We therefore suggest that MHV 3CLpro is not an optimal proxy to study resistance mutations.

**Figure 1:**
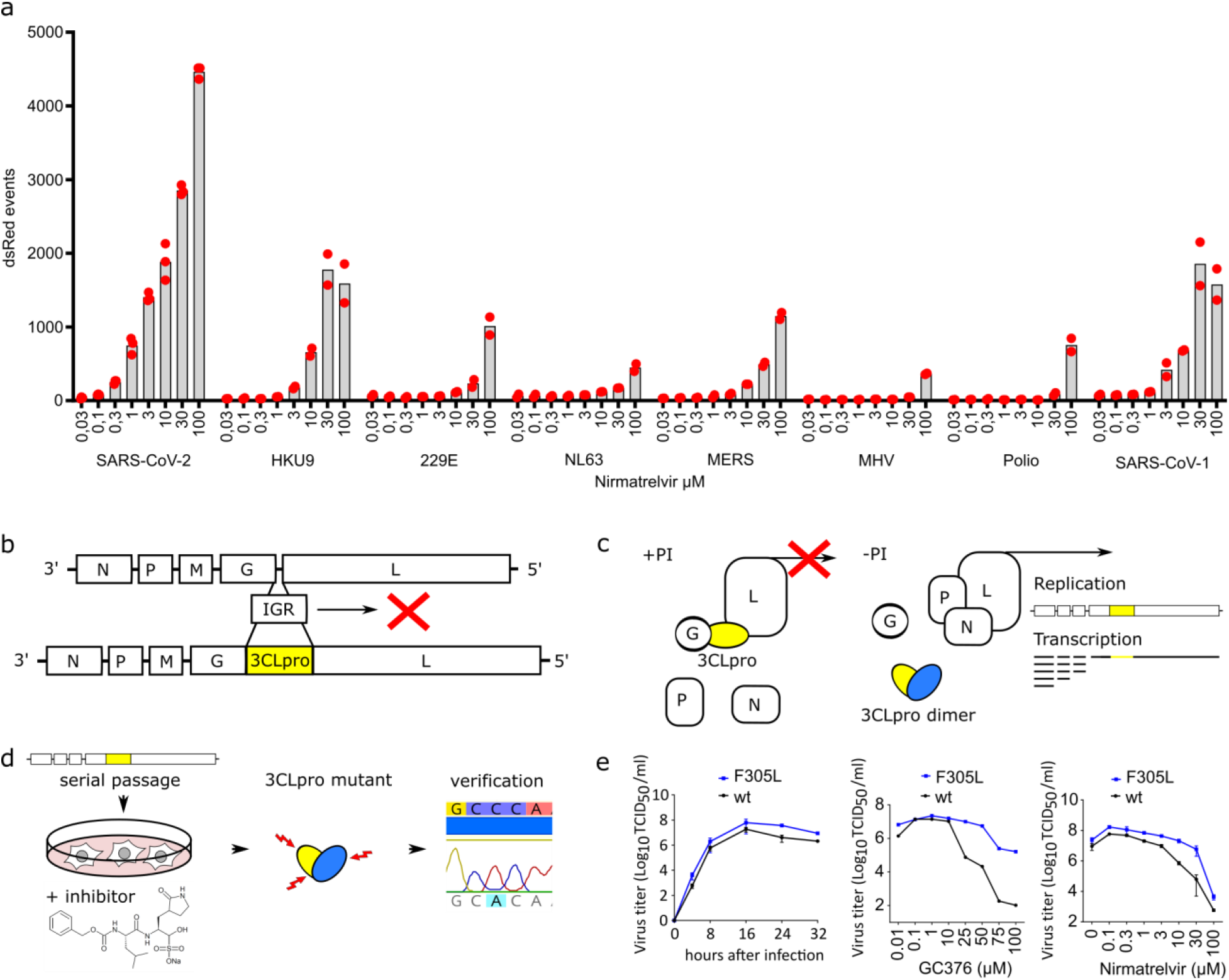
Generation of a VSV-based non-gain-of-function system to predict SARS-CoV-2 3CLpro mutations. **a:** A panel of eight proteases (from SARS-CoV-2, Rousettus bat Coronavirus HKU9, human coronaviruses 229E and NL63, MERS, mouse hepatitis virus (MHV), Poliovirus and SARS-CoV-2) was tested in a gain-of-signal assay. (n of SARS-CoV-2 3CLpro = 3 biologically independent replicates per condition with individual data points shown and average values represented by histogram bars; all other proteases n = 2). **b:** Genomic scheme of VSV with the replacement of the intergenic region (IGR) between the glycoprotein (G) and the polymerase (L) with SARS-CoV-2 3CLpro. **c:** Addition of a protease inhibitor (PI) stalls VSV G, 3CLpro and L in a non-functional polyprotein. In the absence of PI, VSV can replicate and transcribe genes. **d:** Serial passage of VSV-G-3CLpro-L in the presence of a 3CLpro inhibitor leads to resistance mutations. **e:** Replication kinetics and dose responses of wild-type (wt) VSV-G-3CLpro-L and GC376-selected mutant variant with the F305L mutation.

The selection of drug resistant SARS-CoV-2 are considered virus gain-of-function experiments that impose a high medical risk and are thus subject to strict regulations. To generate a safe alternative to gain-of-function studies, we engineered a chimeric VSV variant, where the intergenic region between the glycoprotein (G) and the polymerase (L) was replaced by the 3CLpro of SARS-CoV-2 (**Figure 1b**). Upon translation, G, 3CLpro, and L form a polyprotein, which must be processed by 3CLpro to generate the functional viral proteins G and L. By applying an appropriate protease inhibitor (+PI), this processing is disturbed and therefore viral replication cannot occur (**Figure 1c**). Through passaging the chimeric VSV variant in presence of a protease inhibitor, 3CLpro mutations that are generated by the error-prone viral polymerase (*2, 3*) are selected for resistance to the inhibitor (**Figure 1d**). In a first proof-of-concept study, we selected a mutant against the inhibitor GC376, which acquired the amino acid change in the 3CLpro from phenylalanine to leucine at position 305 (F305L). This virus gained a mildly faster replication kinetic and produced higher titers in the presence of GC376 and nirmatrelvir compared to the parental virus (**Figure 1e**). The F305L mutant was also used for subsequent mutation selection studies to investigate the potential accumulation of multiple mutations in the 3CLpro.

We next used the wild-type and F305L mutant viruses to select nirmatrelvir resistant 3CLpro. BHK-21 in wells of a 96-well plate were infected at a low MOI (0.01). Where cytopathic effect (CPE) was visible in the first passage (25 out of 48 from parental wild-type and 17 out of 48 from parental F305L), supernatants were used for passaging with increasing concentrations (wild-type initial: 30, 2^nd^ 40 and 3^rd^ 50 μM, F305L initial: 50, 2^nd^ 75 and 3^rd^ 100 μM) of nirmatrelvir. We then sequenced the 3CLpro and adjacent parts of G and L in the selected viruses and found 48 mutations within 3CLpro, with viruses carrying from one dominant mutation up to four mutations.

The mutations were distributed over the entire sequence of 3CLpro (**Figure 2, Table S3**). We categorized them into catalytic site and autocleavage site mutants. A third category for all mutations not fitting the first two was chosen as “allosteric” mutants. We searched for those specific mutants in the NCBI Virus data base (*4*) and GISAID EPICOV (*5–7*), and found most of them or at least the same residue with a different mutation in deposited sequences with varying coverage (**Figure 2, Table S3**). We further subdivided GISAID entries into depositions made before and after the emergency use authorization of Paxlovid on 22^th^ December 2022 (**Table S4**).

**Figure 2:**
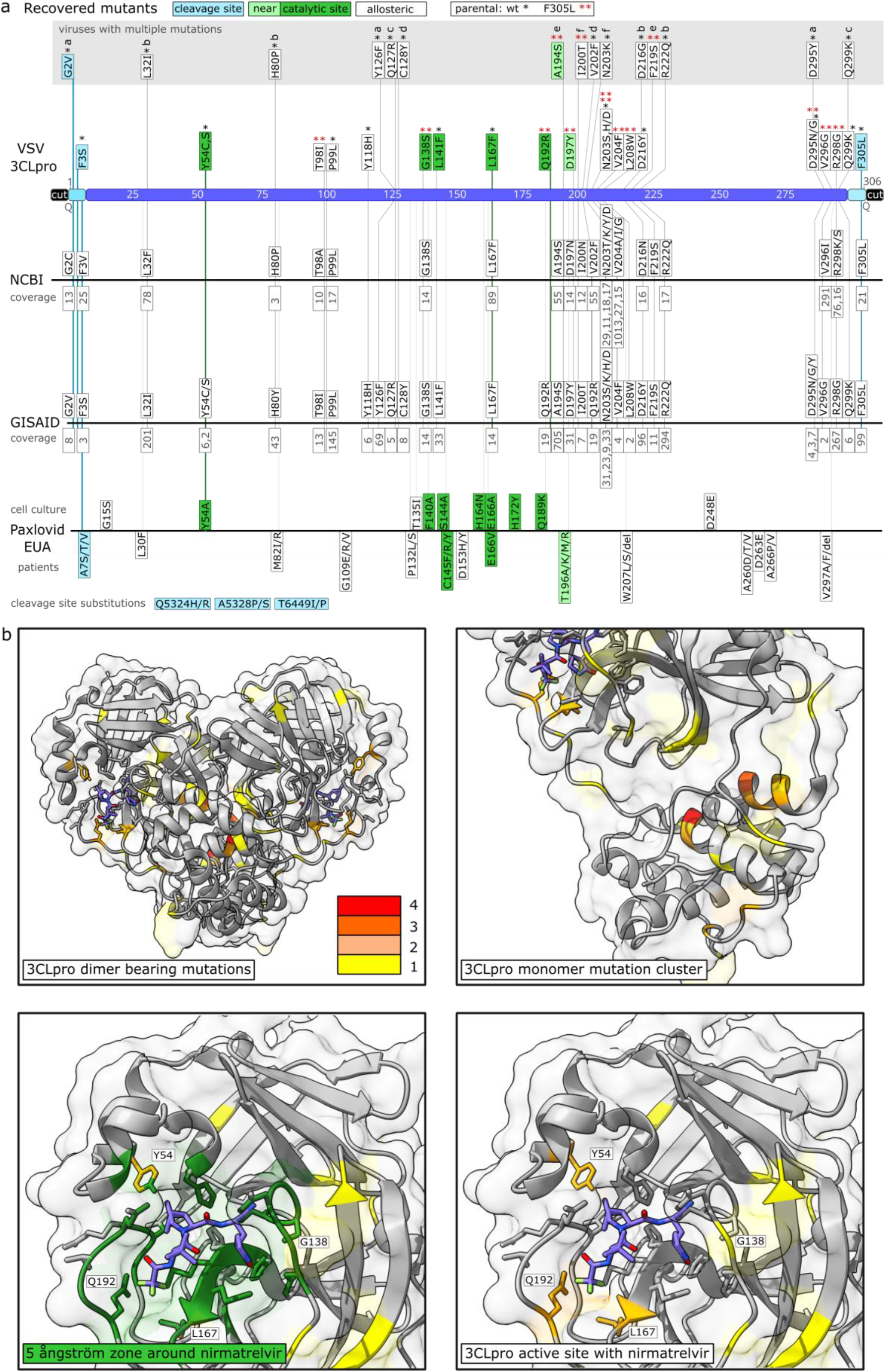
Sequencing of escape mutants and comparison to sequence data bases and Pfizer EUA. **a:** Mutants were recovered from VSV-G-3CLpro-L wild-type (*) and pre-mutated F305L variant (red **). Autocleavage site mutants are colored in turquoise, catalytic site mutants in green, near catalytic site mutants in light green and “allosteric” mutants in white. Viruses with more than one mutation are displayed separately and listed a-f. Frequencies from sequence data bases are displayed below the mutations in grey. Mutations from Pfizer EUA are divided into mutations found in cell culture and such sequenced from treated patients. Additional distant cleavage site mutations outside the 3CLpro open reading frame are depicted at the bottom. The coverage of mutation entries was obtained on march 3^rd^, 2022. **b:** Visualization of mutation-affected residues. Residues that were mutated one time are highlighted in yellow, two times in light orange, three times dark orange and four times in red. The 3CLpro protease dimer with bound nirmatrelvir was visualized in ChimeraX from the Protein Data Bank structure 7vh8. Catalytic center mutations are within a range of 5 ångström as visualized in dark green.

In a recent update of the Paxlovid EUA (18^th^ march), Pfizer included 3CLpro mutants, which they retrieved from patients treated with Paxlovid (**Figure 2a**). They state that it is unclear whether these mutations have clinical relevance (*8*).

To confirm our potential resistance mutations, we chose six virus samples to perform replication kinetics and dose response experiments in comparison with their parental virus variant. We selected viruses for further testing based on two criteria. First, we chose virus variants with catalytic site mutations, because the mechanism of resistance should be straight-forward if steric interactions of catalytic site residues with nirmatrelvir are altered by the mutation. Second, we chose the most abundant mutant outside of the catalytic center. Four samples were derived from wild-type VSV-G-3CLpro-L, two from the pre-mutated F305L variant. The replication kinetics revealed that all variants were still replicating to high titers, suggesting that resistance mutations did not result in a strong negative effect on 3CLpro activity (**Figure 3**). The dose responses showed that wild-type VSV-G-3CLpro-L was inhibited by 10^6^ fold at 100 μM of nirmatrelvir, with an IC_50_ of about 185 nM (**Figure 3a**). We tested two L167F variants, because this mutant arose twice independently. The similarity of their dose responses (**Figure 3b,d)** as well as the low variation of the biological replicates in general suggests that the differences in the level of resistance we observed between the mutants are unlikely artifacts. We observed the strongest resistance phenotype in the double mutant Q192R / F305L, which was almost entirely resistant to nirmatrelvir, replicating to high viral titers (**Figure 3h**) with a pronounced cytopathic effect (**Figure S2**) even in the presence of 100 μM nirmatrelvir.

**Figure 3:**
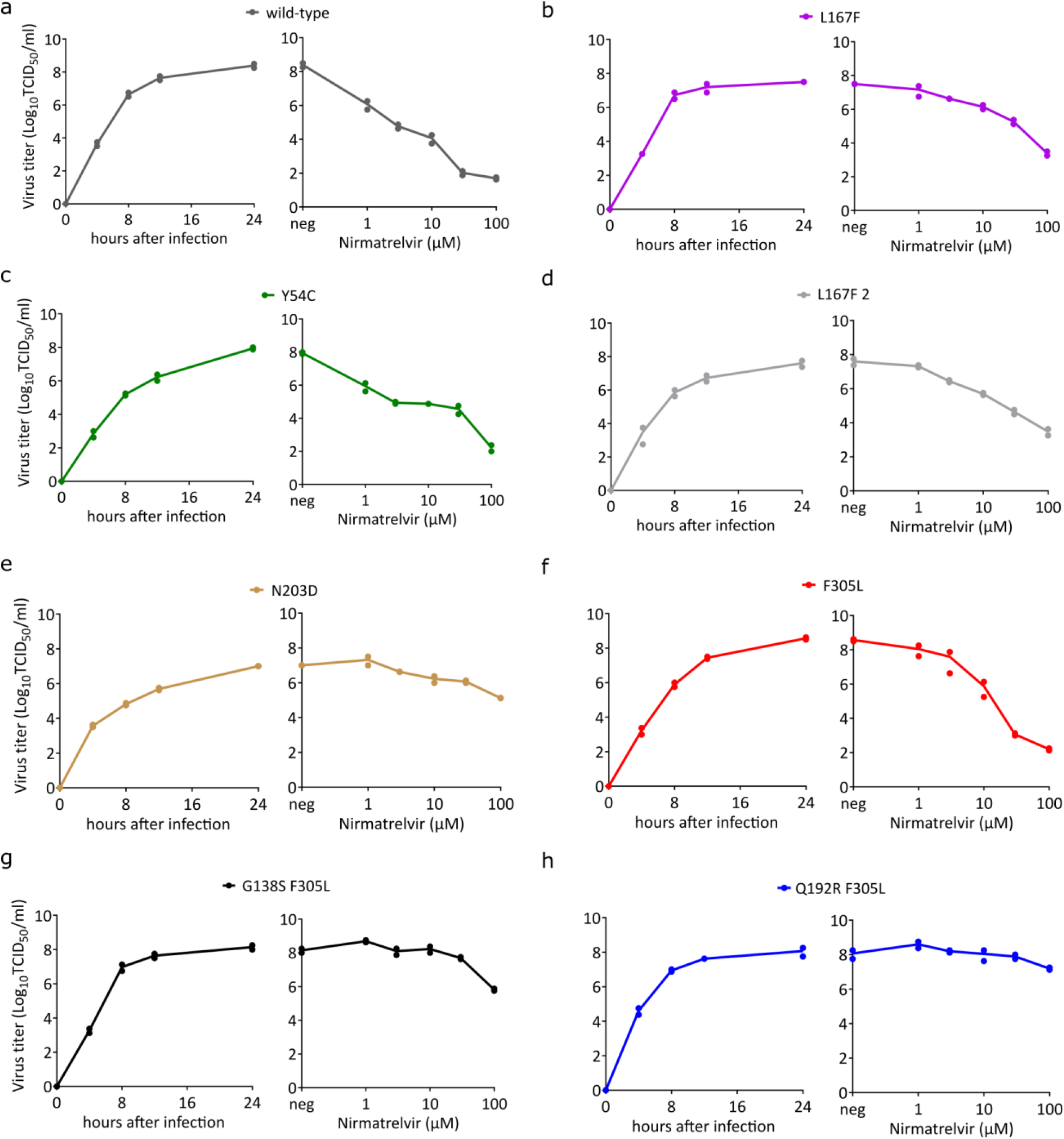
Replication kinetics and nirmatrelvir dose responses of parental VSV-G-3CLpro-L and mutant variants. Replication kinetics and dose responses of wild-type VSV-G-3CLpro-L (**a**), L167F (**b**), Y54C (**c**), L167F-2 (**d**), N203D (**e**), F305L (**f**), G138S/F305L (**g**) and Q192R/F305L (**h**). (n = 2 biologically independent replicates per condition with individual data points shown and connecting lines of mean values)

As shown in **Table S3**, VSV induced 3CLpro mutations occur already in one passage when nirmatrelvir is applied. To strengthen the resistance data of replication-competent VSV-G-3CLpro-L and at the same time exclude the effects of potential additional mutations arising within the dose response experiment, we reintroduced some of the catalytic center mutations into a recently developed protease activity measurement tool based on replication-incompetent VSV (*1*). We found that single catalytic center mutations confer partial resistance against nirmatrelvir (**Figure 4a-d; Figure S3a,b**) and also GC376 (**Figure S3d,e**), which can be further enhanced by introduction of a second mutation in the autocleavage site (**Figure 4e-h**). The mutation Q192R arose in the F305L parental virus. Introducing Q192R alone apparently reduces 3CLpro activity mildly, as we observed by increased values in 3CLpro-On-Q192R at low nirmatrelvir concentrations. Adding F305L as second mutation, thereby restoring the original combination from the double mutant virus, seems to rescue this phenotype (**Figure 4e**). A randomly selected combination of catalytic center mutations led to a strong loss in enzyme activity (**Figure S3g,h**).

**Figure 4:**
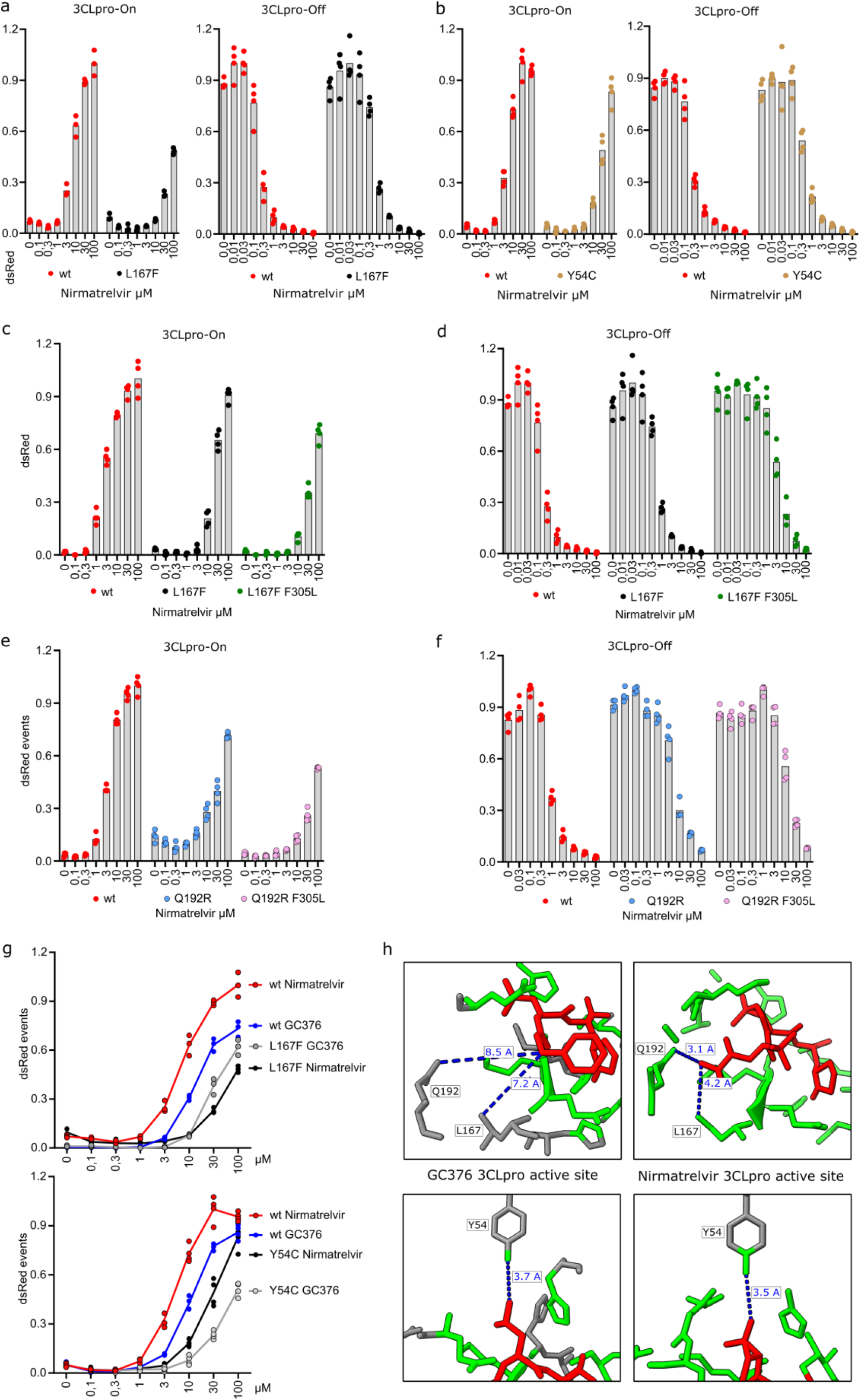
Re-introduction of 3CLpro mutations confirms their resistance phenotype. **a:** Gain-and loss-of-signal assays for the assessment of catalytic site mutation L167F. (3CLpro-On n = 3 biologically independent replicates per condition with individual data points shown and average values represented by histogram bars; 3CLpro-Off n = 4). **b:** Gain-and loss-of-signal assays for the assessment of catalytic site mutation Y54C. (3CLpro-On n = 3 biologically independent replicates per condition with individual data points shown and average values represented by histogram bars; 3CLpro-Off n = 4). **c:** Gain-of-signal assay for the double mutant L167F-F305L compared to single mutant and wild-type 3CLpro. **d:** Loss-of-signal assay for the double mutant L167F-F305L compared to single mutant and wild-type 3CLpro. **e:** Gain-of-signal assays for the of double mutant Q192R-F305L tested with nirmatrelvir. **f:** Loss-of-signal assays for the of double mutant Q192R-F305L tested with nirmatrelvir. **g:** Gain-and loss-of-signal assays for the of single mutants Y54C and L167F tested with GC376 and nirmatrelvir. **h:** GC376 (7k0g) and nirmatrelvir (7vh8) 3CLpro crystal structures with compounds in red and residues in green (within zone of 5 ångström). (**c-f:** n = 4 biologically independent replicates per condition with individual data points shown and average values represented by histogram bars).

Comparing GC376 to nirmatrelvir in the 3CLpro mutants Y54C and L167F mutants directly revealed that mutants react differently to these compounds. Whereas in Y54C GC376 shows a similar decrease as nirmatrelvir, L167F seems to affect GC376 less than nirmatrelvir (**Figure 5a**), despite the higher potency of nirmatrelvir. Leucine 167 is in closer proximity (within 5 ångström) to nirmatrelvir than GC376 (**Figure 5b**), therefore steric hindrance of nirmatrelvir binding through the exchange of leucine with phenylalanine of nirmatrelvir is more likely. We observed a similar effect for the mutant Q192R (**Figure S3c**), where the increase of the EC_50_ was more pronounced with nirmatrelvir than GC376 (**Tables S5,6**). Again, the residue Q192 is closer to nirmatrelvir than GC376 (**Figure 5b**).

Especially in the 3CLpro-On construct, nirmatrelvir EC_50_s are high. We sought to improve the assays sensitivity by changing the read-out method from a FluoroSpot to a FACS reading. With this approach we could decrease the EC_50_ of the wild-type 3CLpro-On to 0.91 μmol of nirmatrelvir (**Figure S3f, Table 4**).

## Discussion

More than two years into the SARS-CoV-2 pandemic, several vaccines, a protease inhibitor (Nirmatrelvir) and a polymerase inhibitor (Molnupiravir) are deployed against this virus. Nirmatrelvir, a specific and potent protease inhibitor for SARS-CoV-2 3CLpro was developed by Pfizer and licensed as combination drug with ritonavir to prolong the half-life of nirmatrelvir, thereby creating Paxlovid.

However, new variants have circumvented current vaccines and are likely to do so for the novel antiviral drugs. In Pfizer’s initial resistance studies leading to the emergency use authorization, the 3CLpro of a related coronavirus, mouse hepatitis virus (MHV), was utilized to generate resistance mutants. The 3CLpro of SARS-CoV-2 and MHV share 50 % sequence identity. In this study we first tested different proteases from close and distantly related viruses, among others the 3CLpro of MHV. We observed that MHV 3CLpro responds only mildly to nirmatrelvir in our recently developed gain-of-signal assay (*1*). Although the structures of SARS-CoV-2 and MHV 3CLpro are conserved well, we propose that the low amino acid sequence identity alters the binding pocket affinity to nirmatrelvir sufficiently to generate a certain resistance against the inhibitor. Key corresponding residues of the binding pocket (within 5 ångström or less) are different, namely H164 - Q, M165 - L, P168 - S, V186 - R, R188 - A, T190 - V and A191 - V. Furthermore, amino acid changes that occurred in our mutation induction assay, Y126F and F305L are already present in the MHV 3CLpro sequence. Taken together we argue that MHV 3CLpro was not an optimal proxy for resistance studies.

In our study, we predict mutations against the compound nirmatrelvir with a safe, non-gain-of-function system based on the vesicular stomatitis virus (VSV). Recently, chimeric VSV variants with SARS-CoV-2 spike were used to predict spike protein mutations by selecting against neutralizing sera (*9–11*). The fast occurrence of mutations was facilitated in those studies by the high error rate of the VSV polymerase (*3*). Here, we show that this high error rate leads to 3CLpro mutations that confer resistance against nirmatrelvir. To this end, we leverage a surrogate polyprotein consisting of the viral glycoprotein, 3CLpro and its polymerase built into VSV, which renders the virus dependent on 3CLpro activity for its replication. In a small proof-of-concept study with the inhibitor GC376 we retrieve the mutant F305L, which displays a slightly faster replication kinetic as well as resistance against GC376 and nirmatrelvir. These properties could be explained by the optimization of the C-terminal autocleavage site from FQ (P_2_/P_1_) to LQ (P_2_/P_1_), which is a preferred recognition motif (*12, 13*). We then test both wild-type and the pre-mutated F305L against nirmatrelvir to study if additional mutations can accumulate in 3CLpro, which will likely occur in nature as precedented by multi-mutant spike variants such as Omikron. By applying a selection pressure with nirmatrelvir, mutations from both wild-type and F305L were selected that ultimately allow the mutants to escape the inhibitor. We confirmed their resistance phenotype by dose response experiments and re-introduction of mutations into recently developed protease activity measurement systems (*1*). As we show by the example of GC376 and nirmatrelvir, mutants react differently to these compounds due to their structural characteristics. This observation could aid future guided compound structure adaptation.

One technical shortcoming of this study, especially in the 3CLpro-On construct, is that the nirmatrelvir EC_50_s are high. The screening method used in this study to assess mutants was originally developed as high-throughput screening tool for 3CLpro inhibitors (*1*). Using a FluoroSpot reader allows fast sampling of a large number of wells within minutes, which is essential for high-throughput screening efforts. Sampling speed however means that this particular readers signal threshold sensitivity is low. We improved the assays sensitivity by changing the read-out method from a FluoroSpot to FACS analysis. FACS analysis, although being more time consuming, proved to be more sensitive to low-level fluorescent cells, whereas FluoroSpot cell counting relied on highly fluorescent cells to clear the detection threshold. Therefore, FACS read-out also captured milder inhibition and resulted in a more gradual signal increase and therefore lower EC_50_s. With this approach we could decrease the EC_50_ of the wild-type 3CLpro-On 0.91 μM of nirmatrelvir, which is closer to the published range of 74.5 (66.5 – 83.4) nM (*14*). A further limitation is that the effect of the mutations on replication fitness of SARS-CoV-2 remains unclear. Although the VSV-chimeric viruses containing resistant 3CLpro showed no fitness loss, this needs to be confirmed for SARS-CoV-2 containing the identified 3CLpro mutations. Along this line, the degree of resistance to the protease inhibitors conferred by these mutations also needs to be studied further in the context of SARS-CoV-2.

In this study, we identified several mutations such as Y54C, G138S, L167F, Q192R, A194S andF305L in the SARS-CoV-2 3CLpro that confer resistance to the protease inhibitor nirmatrelvir. These findings argue for a highly selective application of protease inhibitors to risk patients, as extensive unselective use is expected to rapidly lead to emergence of drug resistance.

## Methods

### Cloning strategies

The chimeric VSV variant with 3CLpro instead of the intergenic region between G and L was cloned via Gibson assembly (NEB, Ipswich, USA) (*15*). A VSV-G plasmid (*16*) was digested with KpnI and HpaI (NEB), removing a C-terminal part of G, the intergenic region and a small N-terminal part of L. Insert fragments were generated as follows. Missing C-terminal G with an additional overhang to the N-terminal cleavage site of 3CLpro was amplified with primers 33n-before-KpnI-for and G-cut1-rev. 3CLpro with its N-and C-terminal cleavage sites and a C-terminal overhang to L was amplified from Wuhan-1 (NCBI Reference Sequence: NC_045512.2) cDNA with primers cut1-for and cut2-L-rev. The N-terminal missing L sequence was amplified with primers cut2-L-for and 33n-after-HpaI-rev. For subsequent Gibson assembly, the fragments were ligated in a fusion PCR using the outer primers 33n-before-KpnI-for and 33n-after-HpaI-rev with all three fragments as templates. The construct sequence was submitted to GenBank (VSV-G-3CLpro-L: 2568407).

**Table S1:**
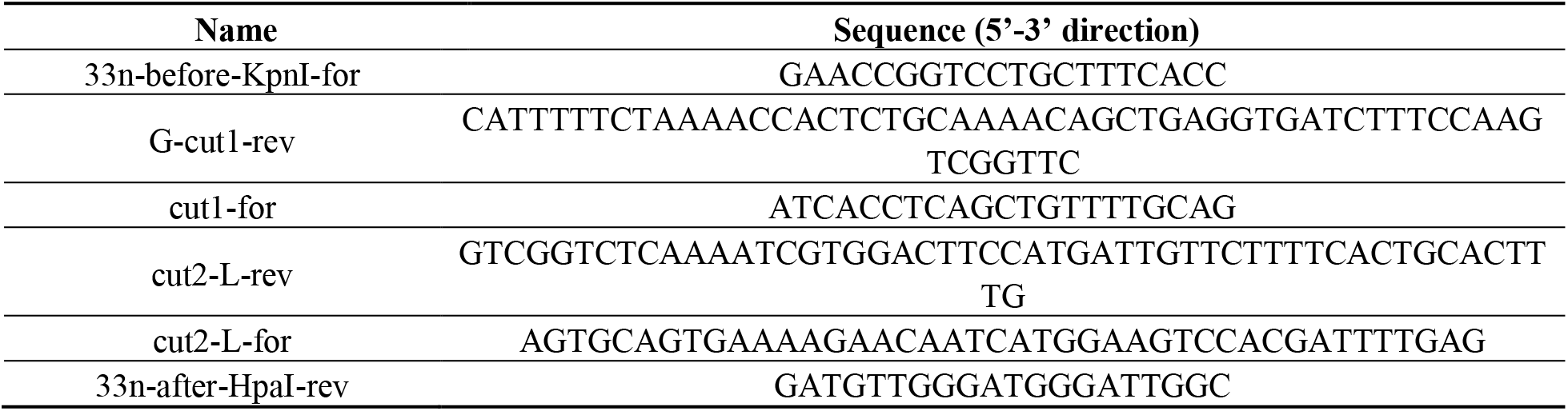
cloning primer for VSV vectors.

3CLpro-Off and-On point mutants were generated by mutagenic Gibson assembly on parental plasmids (GenBank accession codes: 3CLpro-Off: 25684003; 3CLpro-On: 2568399). For 3CLpro-Off mutants, a lentiviral expression plasmid expressing VSV L (identical sequence as blasticidin 3CLpro-Off plasmid without GFP and 3CLpro) was digested with HpaI, which removed the cPPT/CTS and CMV promoter sequences and a small N-terminal part of L. This missing sequence was replaced with the identical sequence from 3CLpro-Off with the addition of the N-terminal 3CLpro sequence up to the respective mutation site with primers blasticidin-for and 3CLpro-*mut-x*-rev. The C-terminal part of 3CLpro and the small missing fragment of L were generated by PCRs on parental vectors with primers 3CLpro-*mut-x*-for and 33n-after-HpaI-rev.

For 3CLpro-On mutants, a lentiviral hygromycin vector (modified from Addgene pLenti CMVie-IRES-BlastR accession: #119863) was digested with NheI and PacI. N-terminal 3CLpro insert fragments with vector overhangs were generated with hygro-P-for and 3CLpro-*mut-x*-rev. C-terminal 3CLpro insert fragments with vector overhangs were generated with 3CLpro-*mut-x*-for and P-hygro-rev. Double mutants were cloned by repeating the site directed mutagenesis with a second primer pair in combination with Gibson assembly on an already mutant-bearing plasmid.

**Table 2:**
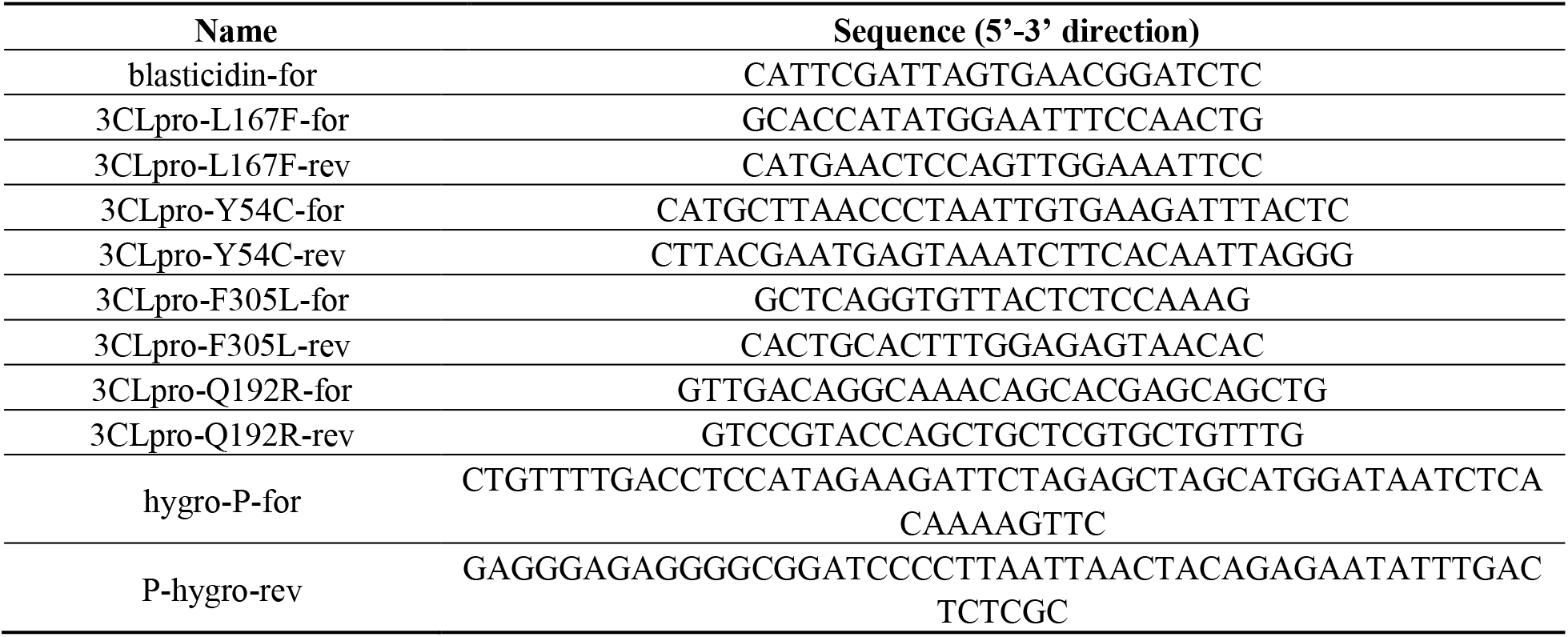
cloning primer for 3CLpro-Off and-On mutant variants.

### Cell lines

BHK-21 cells (American Type Culture Collection, Manassas, VA) were cultured in Glasgow minimum essential medium (GMEM) (Lonza, Basel, Switzerland) supplemented with 10% fetal calf serum (FCS), 5% tryptose phosphate broth, 100 units/ml penicillin and 0.1 mg/ml streptomycin (P/S) (Gibco, Carlsbad, California, USA). 293T cells (293tsA1609neo), and 293-VSV (293 expressing N, P-GFP and L of VSV) (*17*) and U87 cells (ATCC) were cultured in Dulbecco’s Modified Eagle Medium (DMEM) supplemented with 10% FCS, P/S, 2% glutamine, 1x sodium pyruvate and 1x non-essential amino acids (Gibco).

### Virus recovery

VSV-G-3CLpro-L was rescued in 293T cells by CaPO_4_ transfection of whole-genome VSV plasmids together with T7-polymerase, N-, P-, M-, G-and L expression plasmids as helper plasmids (*18*). Briefly, genome and helper plasmids are transfected into 293T in the presence of 10 μM chloroquine to avoid lysosomal DNA degradation. After 6-16 hours, chloroquine is removed and cells cultured until cytopathic effects occur. M and G proteins as helper plasmids are optional in the recovery of VSV, but were chosen here as a precaution to support the rescue of a potentially attenuated virus variant. After the rescue, viruses were passaged on 293-VSV cells and plaque purified twice on BHK-21 cells. ΔP and ΔL VSV variants expressing dsRed were produced on replication supporting 293-VSV cells. VSV-G-3CLpro-L was fully replication competent and therefore produced on BHK-21 cells.

### Multi-step growth curve, TCID50 assay and dose responses

For multi-step growth kinetics, 10^5^ BHK-21 cells per well were seeded in 24-well plates one day before infection. Cells were infected in duplicates with a multiplicity of infection (MOI) of 0.5 of VSV 3CLpro wild-type or mutant variants. One hour after infection the medium was removed, cells were washed with PBS, and fresh medium was added. Supernatant was collected at the indicated time points and stored at −80 °C until further analysis. For quantification, the 50% tissue culture infective dose (TCID_50_) assay was performed as described previously (*19*). In short, 100 μl of serial dilutions of virus were added in octuplicates to 10^3^ BHK-21 cells seeded in a 96-well plate. Six days after infection the TCID_50_ were read out and titers were calculated according to the Kaerber method (*20*).

For initial dose response experiments, 5 × 10^4^ BHK-21 cells per well were seeded in 48-well plates one day before infection. Cells were infected in duplicates with a MOI of 0.05 of VSV 3CLpro wild-type or mutant variants and indicated concentrations of nirmatrelvir were added to the wells. After 48 hours, supernatants were collected and titrated with TCID_50_.

For mutant comparing dose response experiments, 5 x 10^4^ BHK-21 cells per well were seeded in 48-well plates one day before infection. Cells were infected in duplicates with a MOI of 0.05 of VSV 3CLpro wild-type or mutant variants and indicated concentrations of nirmatrelvir added to the wells. To prevent initial escape or further mutation in wild-type or already mutation-bearing viruses, respectively, supernatants of all viruses were collected after the first mutant (Q192R, F305L) showed massive cytopathic effect at 100 μM nirmatrelvir (ca. 24 hours after infection).

### Viral RNA isolation and 3CLpro sequencing

VSV-G-3CLpro-L RNA was isolated with E.Z.N.A.^®^ Viral RNA Kit (Omega Bio-Tek Inc., Norcross, Georgia, USA) or NucleoSpin^®^ RNA Virus (Macherey-Nagel GmbH, Düren, Germany). BHK-21 cells were infected with VSV-G-3CLpro-L wild type and F305L (3CLpro) mutant in 96-well plates. Virus-containing supernatants were collected and the RNAs purified according to manufacturers’ instructions. Then, cDNA was synthesized from isolated viral RNA via RevertAid RT Reverse Transcription Kit (Thermo Scientific^TM^ Waltham, Massachusetts). 3CLpro sequence was amplified by PCRs with primers for: CTCAGGTGTTCGAACATCCTCAC and rev: GATGTTGGGATGGGATTGGC and sent for sequencing (MicroSynth AG, Balgach, Switzerland). Obtained sequences were mapped to the 3CLpro-wt (Wuhan-1) reference sequence in Geneious Prime 2022.0.2 and examined for mutations.

### Mutation induction assay

10^4^ BHK-21 cells per well in a 96-well plate were seeded one day before infection with wild-type VSV-G-3CLpro-L or pre-mutated VSV-G-3CLpro-L-F305L at an MOI of 0.01 and indicated nirmatrelvir doses. Each virus variant occupied 48 wells of the 96-well plate. Wells that displayed cytopathic effect after two days (25 out of 48 from parental wild-type and 17 out of 48 from parental F305L) were further passaged with increasing concentrations of nirmatrelvir with each passage (wild-type: 30, 40 and 50 μM, F305L: 50, 75 and 100 μM). **Table S3** indicates at which passage a pure mutant virus could be distinguished via Sanger sequencing, i.e. only one base-pair peak appeared in the chromatogram instead of a mixture with the parental virus. Only pure mutants are displayed in **Figure 2** and **Table S3**.

### Screening assay with fluorospot read-out

3×10^5^ cells were seeded in 6-well plates and transfected with 3CLpro plasmids via TransIT-PRO (Mirus Bio LLC, Madison, WI, USA) and incubated overnight. Then, cells were seeded into a 96-well plate with fifteen thousand cells per well (cell number can be adjusted up to 20.000 cells per well for toxic compounds). Directly after seeding, compounds and virus (MOI: 0.1) were added to wells. After 48 hours, supernatants were removed, and fluorescent spots counted in a Fluoro/ImmunoSpot counter (CTL Europe GmbH, Bonn, Germany) with the manufacturer-provided software CTL switchboard 2.7.2. 90 % of each well area was scanned concentrically to exclude reflection from the well edges and counts were normalized to the full area. Automatic fiber exclusion was applied while scanning. The excitation wave length for RFP was 570 nm, the D_F_R triple band filter was used to collect fluorescence. Manual quality control for residual fibers was also performed. To increase comparability between 3CLpro-On and-Off signals, we normalized dsRed events with the following strategies. In 3CLpro-On, the highest signals (i.e. high compound concentrations) would not reach the same levels. Therefore, we normalized to the highest mean of the experiment, i.e. the wild-type signal. In 3CLpro-Off, untreated wells reached the same signal yield in wild-type and mutants. Therefore, we normalized the signal to each individual highest mean of the construct.

### Screening assay with fluorescence-activated cell scanning read-out

3×10^5^cells were seeded in 6-well plates and transfected with 3CLpro plasmids via TransIT-PRO (Mirus Bio LLC, Madison, WI, USA) and incubated overnight. Then, cells were seeded into a 96-well plate with 1.5×10^4^ cells per well (cell number can be adjusted up to 2×10^4^ cells per well for toxic compounds). Compound and virus (MOI 0.1) were added in 50 μl to reach desired concentrations. After two days, cells were detached with 0.05 % Trypsin-EDTA (Gibco) and transferred to a 96-well round-bottom plate (TPP Techno Plastic Products AG, Switzerland) for automatic sampling by fluorescence-activated cell scanning (BD FACSCanto II).

### Half-maximal effective/inhibiting concentrations (EC_50_ and IC_50_)

EC_50_ and IC_50_ calculations and statistical analysis were performed with GraphPad Prism 9. Raw data of 3CLpro-On and 3CLpro-Off dose response experiments were normalized using the built-in normalization function of Prism. To perform the calculations, we used two different non-linear regression analyses: [Agonist] vs. normalized response and [Inhibitor] vs. normalized response for EC50 (3CLpro-On) and IC50 (3CLpro-Off), respectively.

**Figure S1:**
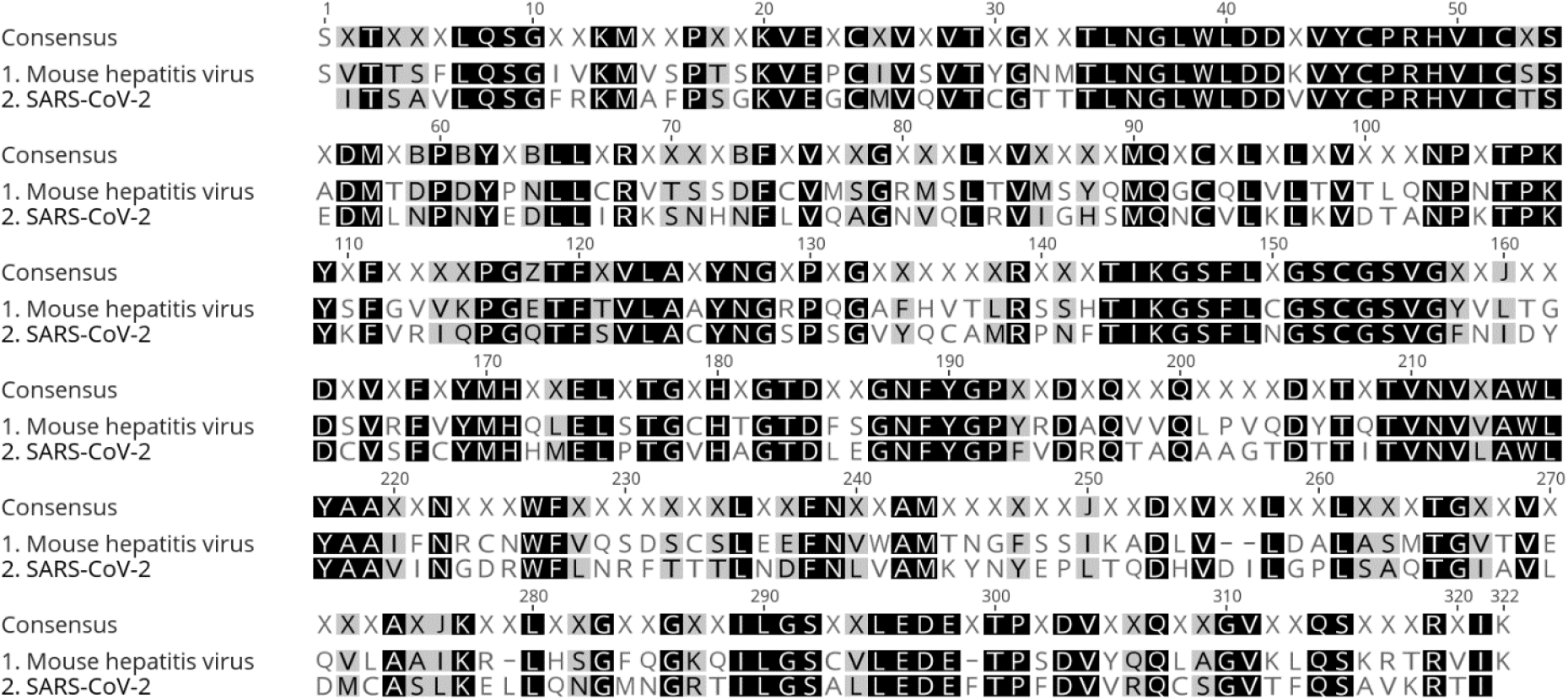
Muscle sequence alignment of SARS-CoV-2 and mouse hepatitis virus 3CLpro shows 50% identity of amino acid sequences.

**Table S3:** 3CLpro mutations derived from VSV-G-3CLpro-L virus.

| Passage | Parental | Amino acid change | Localization | NCBI (coverage 3<sup>rd</sup> march 2022) | GISAID (coverage 3<sup>rd</sup> march 2022) |
| --- | --- | --- | --- | --- | --- |
| 1 | wt | D216Y | allosteric | D216N (16) | D216Y (96) |
| 1 | wt | G2V, D295Y | cleavage site, allosteric | G2C (13) | G2V (8), D295Y (7) |
| 1 | F305L | F3S | cleavage site | F3V (25) | F3S (3) |
| 1 | wt | L32I, H80P, D216G | allosteric, allosteric, allosteric | L32F (78), H80Y (43) | L32I (201), H80P (3), D216G (11) |
| 1 | wt | L167F | catalytic site | L167F (89) | L167F (14) |
| 1 | wt | V204F | allosteric | V204A (1013), V204I (27), V204G (15) | V204F (4) |
| 1 | wt | Y54C | catalytic site |  | Y54C (6) |
| 1 | F305L | Y54S | catalytic site |  | Y54S (2) |
| 1 | wt | Q299P | cleavage site |  |  |
| 1 | wt | Y118H | allosteric |  | Y118H (6) |
| 2 | F305L | A194S, F219S | near catalytic site, allosteric | A194S (55) | A194S (705), F219S (11) |
| 2 | F305L | D197Y | near catalytic site | D197N (14) | D197Y (31) |
| 2 | F305L | D295G | cleavage site |  | D295G (3) |
| 2 | wt | D295N | cleavage site |  | D295N (4) |
| 2 | F305L | F219S | allosteric |  | F219S (11) |
| 2 | F305L | G138S | catalytic site | G138S (14) | G138S (14) |
| 2 | wt | L208W | allosteric |  | L208W (2) |
| 2,3 | wt | N203D | allosteric | N203D (17) | N203D (33) |
| 2 | F305L | N203H | allosteric | N203Y (18) | N203H (9) |
| 2 | F305L | N203K | allosteric | N203K (11) | N203K (23) |
| 2 | wt | N203S | allosteric | N203T (29) | N203S (31) |
| 2 | F305L | Q192R | catalytic site |  | Q192R (19) |
| 2 | wt | Q299K | cleavage site |  | Q299K (6) |
| 2 | F305L | R298G | cleavage site | R298K (76), R298S (16) | R298G (267) |
| 2 | wt | C128Y, V202F | allosteric |  | C128Y (8), V202F (32) |
| 2 | F305L | V296G | cleavage site | V296I (291) | V296G (2) |
| 3 | F305L | T98I | allosteric | T98A (10) | T98I (13) |
| 3 | wt | P99L | allosteric | P99L (17) | P99L (145) |
| 3 | wt | Q127R, Q299K | allosteric, cleavage site |  | Q127R (5), Q299K (6) |
| 3 | wt | F305L | cleavage site | F305L (21) | F305L (99) |
| 3 | F305L | I200T, N203K | allosteric | I200N (12), N203K (11) | I200T (7), N203K (23) |
| 3 | F305L | V204F | allosteric |  | V204F (4) |
| 3 | wt | G2V, Y126F,<br>D295Y | cleavage site,<br>allosteric,<br>allosteric | G2C (13) | G2V (8), Y126F<br>(69), D295Y (7) |
| 3 | wt | D216Y | allosteric |  | D216Y (96) |
| 3 | wt | L141F | catalytic site |  | L141F (33) |
| 3 | wt | L32I, H80P,<br>D216G,<br>R222Q | allosteric,<br>allosteric,<br>allosteric,<br>allosteric | L32F (78), H80Y (43),<br>R222Q (17) | L32I (201), H80P<br>(3), D216G (11),<br>R222Q (294) |

**Table S4:**
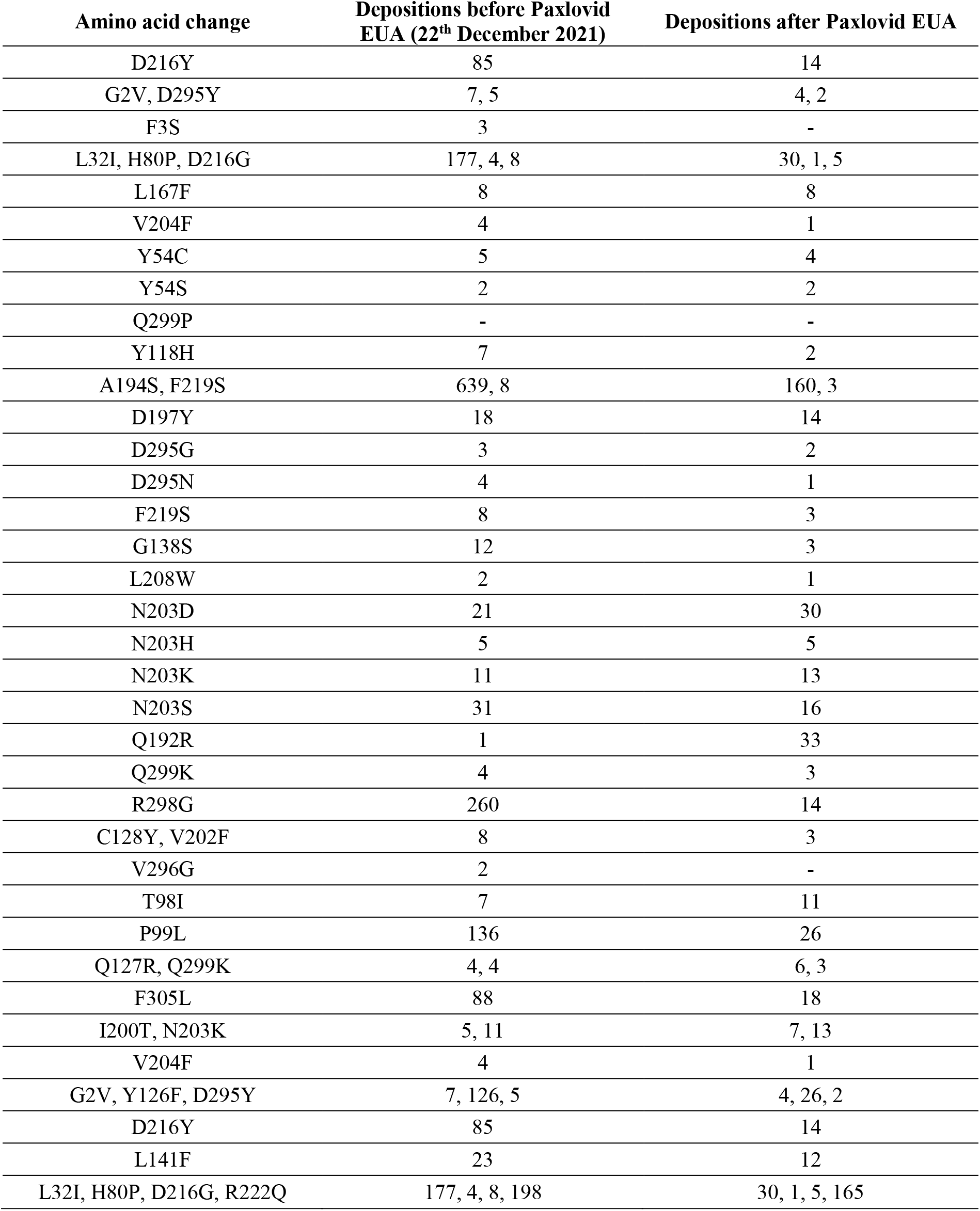
Deposition count in GISAID data base of 3CLpro mutations derived from VSV-G-3CLpro-L virus before and after 22^th^ December 2021 (retrieval 1^st^ May 2022).

**Figure S2:**
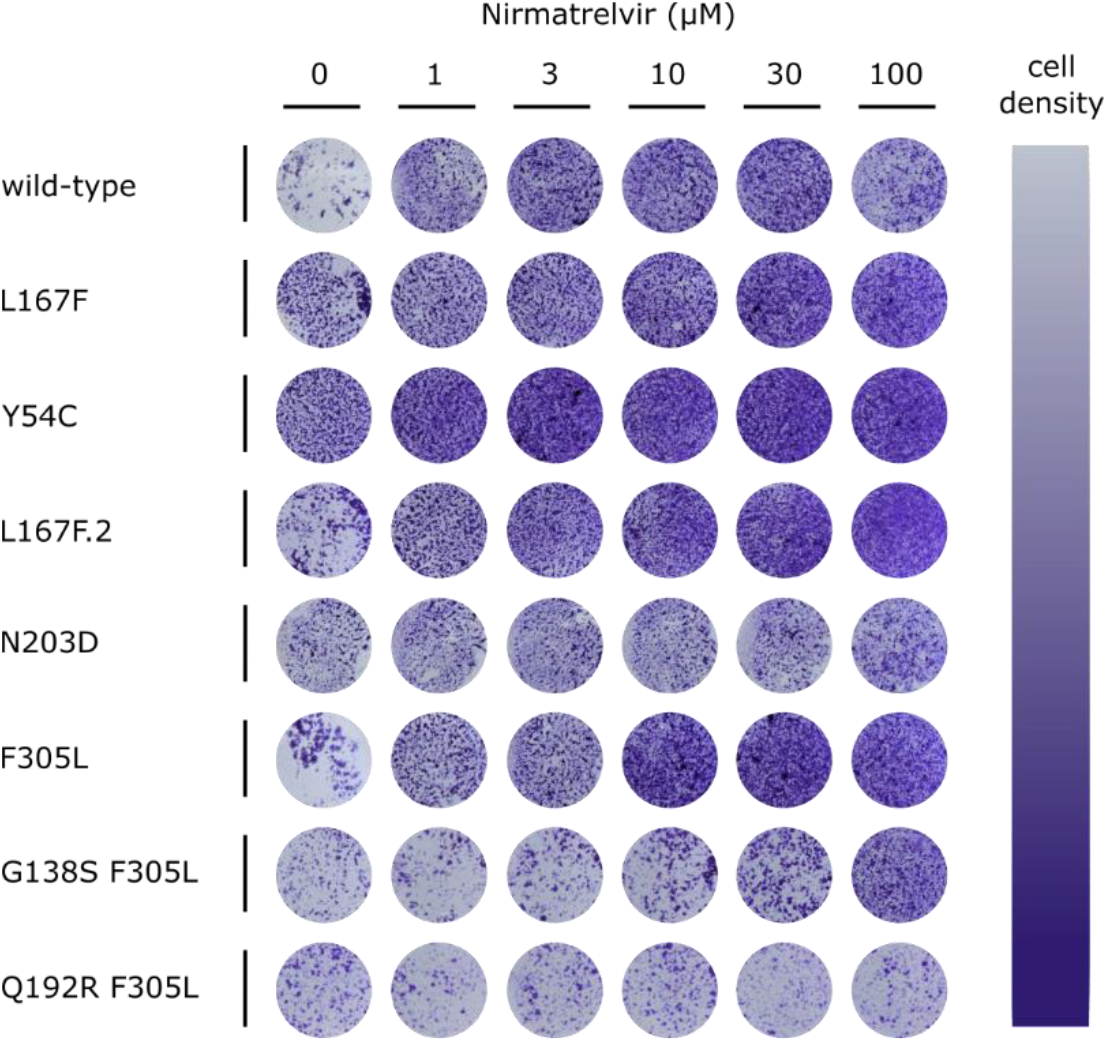
Crystal violet staining of dose response experiments. Cells were stained with crystal violet after removal of supernatant for titration. Lower cell density indicates cytopathic effect.

**Figure S3:**
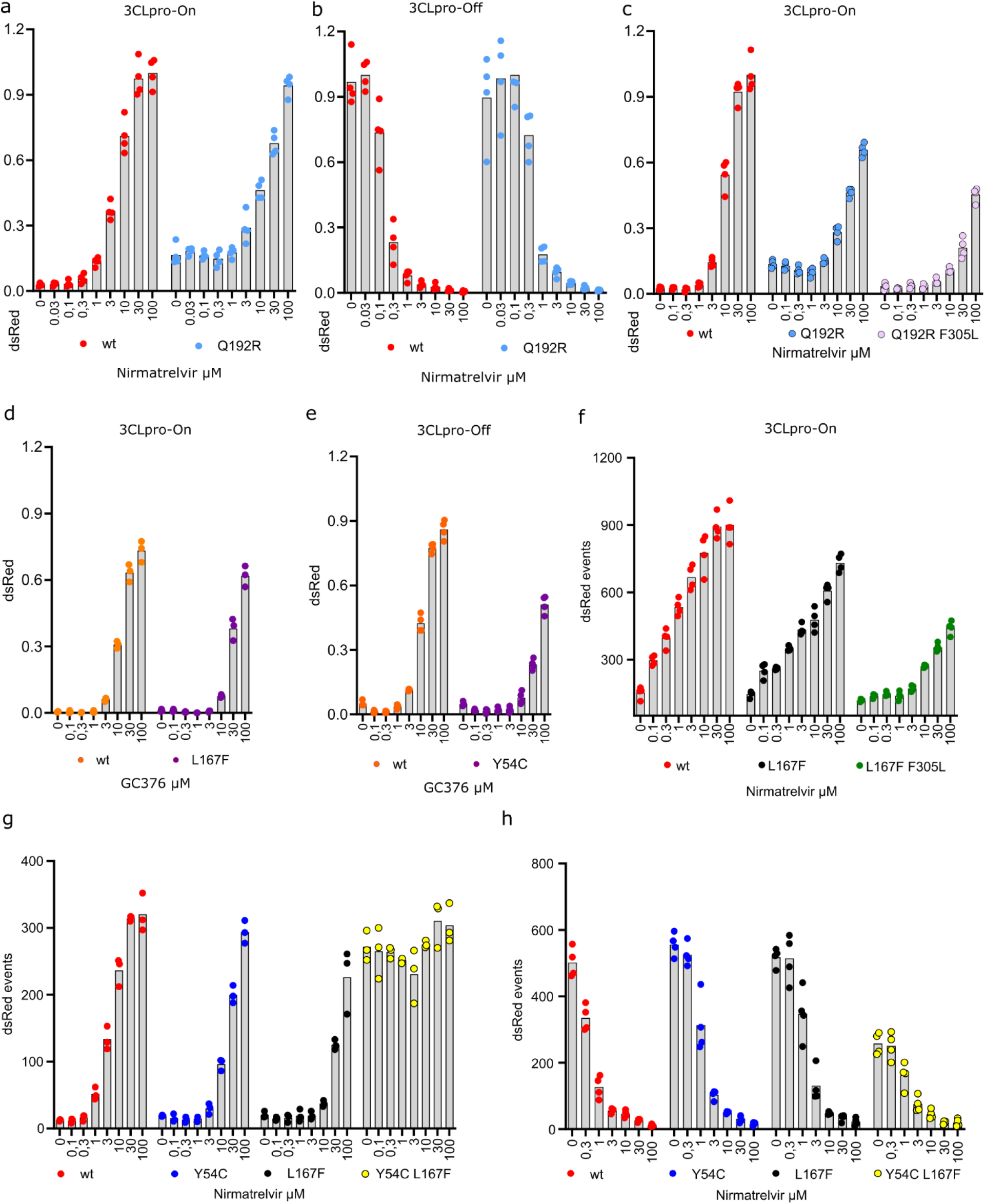
Figure S3: Supporting data for re-introduced mutations into 3CLpro-On and-Off. **a:** 3CLpro-On wild-type vs. Q192R mutant. b: 3CLpro-On wild-type vs. Q192R mutant. (n=4 biologically independent replicates per condition with average values represented by histogram bars) c: 3CLpro-On wild-type vs. L167F mutant treated with GC376. d: 3CLpro-On wild-type vs. Y54C mutant treated with GC376. (n=3 biologically independent replicates per condition with average values represented by histogram bars). e: Dose response experiment from Figure 3c read-out with FACS. Values were not normalized. (n=4 biologically independent replicates per condition with average values represented by histogram bars). f: 3CLpro-On wild-type vs. Y54C, L167F and Y54C-L167F mutants. Values were not normalized. (n=3 biologically independent replicates per condition with average values represented by histogram bars). g: 3CLpro-Off wild-type vs. Y54C, L167F and Y54C-L167F mutants. Values were not normalized. (n=4 biologically independent replicates per condition with average values represented by histogram bars).

**Table S5:** Nirmatrelvir EC_50_ values of 3CLpro-On and-Off wild-type and mutant variants

| Wild-type vs L167F vs L167F+F305L |  |  |  |  |
| --- | --- | --- | --- | --- |
|  | 3CLpro-On<br>EC<sub>50</sub> (CI 95%) | EC<sub>50</sub> fold change<br>(vs wt) | 3CLpro-Off<br>IC<sub>50</sub> (CI 95%) | IC<sub>50</sub> fold change<br>(vs wt) |
| Wild type | 2,89<br>(2,53 – 3,32) | 1 | 0,196<br>(0,16 – 0,25) | 1 |
| L167F | 18,67<br>(14,81 – 23,55) | ~6,4 | 0,59<br>(0,45 – 0,72) | ~3 |
| L167F+F305L | 26,74<br>(21,09 – 33,97) | ~9,2 | 3,32<br>(2,69 – 4,09) | ~16,9 |
| Wild-type vs Y54C vs L167F+Y54C |  |  |  |  |
|  | 3CLpro-On<br>EC<sub>50</sub> (CI 95%) | EC<sub>50</sub> fold change<br>(vs wt) | 3CLpro-Off<br>IC<sub>50</sub> (CI 95%) | IC<sub>50</sub> fold change<br>(vs wt) |
| Wild type | 4,23<br>(3,49 – 5,11) | 1 | 0,44<br>(0,36 – 0,52) | 1 |
| Y54C | 17,76<br>(14,27 – 22,13) | ~4,2 | 1,16<br>(0,91 – 1,48) | ~2,7 |
| L167F | 27,94<br>(19,71 to 39,79) | ~6,6 | 1,5<br>(1,12 – 2) | 3,44 (~3,4) |
| L167F+Y54C | \ | \ | 1,42<br>(1,05 – 1,93) | 3,27 (~3,3) |
| Wild-type vs Q192R vs Q192R+F305L |  |  |  |  |
|  | 3CLpro-On<br>EC<sub>50</sub> (CI 95%) | EC<sub>50</sub> fold change<br>(vs wt) | 3CLpro-Off<br>IC<sub>50</sub> (CI 95%) | IC<sub>50</sub> fold change<br>(vs wt) |
| Wild type | 4,03<br>(3,44 – 4,73) | 1 | 0,78<br>(0,6 – 1) | 1 |
| Q192R | 20,80<br>(16,45 – 26,29) | ~5,2 | 4,61<br>(3,86 – 5,51) | ~5,9 |
| Q192R+F305L | 26,46<br>(21,16 – 33,11) | ~6,6 | 9,702<br>(7,15 – 13,15) | ~12,5 |
| Wild-type vs L167F vs L167F+F305L (with FACS) |  |  |  |  |
|  | 3CLpro-On<br>EC<sub>50</sub> (CI 95%) | EC<sub>50</sub> fold change<br>(vs wt) |  |  |
| Wild type | 0,91<br>(0,7 – 1,19) | 1 |  |  |
| Q192R | 3,53<br>(2,39 – 5,22) | ~3,8 |  |  |
| L167F+F305L | 11,07<br>(9,35 – 13,10) | ~12,5 |  |  |

**Table S6:** GC376 EC_50_ values of 3CLpro-On and-Off wild-type and mutant variants

| Wild-type vs L167F vs L167F+F305L |  |  |
| --- | --- | --- |
|  | 3CLpro-On<br>EC<sub>50</sub> (CI 95%) | EC<sub>50</sub> fold change<br>(vs wt) |
| Wild type | 12,35<br>(2,53 – 3,32) | 1 |
| L167F | 24,23<br>(17,90 – 32,89) | ~2 |
| L167F+F305L | 45,51<br>(31,99 – 65,35) | ~3,7 |
| Wild-type vs Y54C |  |  |
|  | 3CLpro-On<br>EC<sub>50</sub> (CI 95%) | EC<sub>50</sub> fold change<br>(vs wt) |
| Wild type | 9,63<br>(7,88 – 11,77) | 1 |
| Y54C | 30,78<br>(23,58 – 40,29) | ~3,2 |
| Wild-type vs Q192R vs Q192R+F305L |  |  |
|  | 3CLpro-On<br>EC<sub>50</sub> (CI 95%) | EC<sub>50</sub> fold change<br>(vs wt) |
| Wild type | 8,82<br>(7,093 – 10,96) | 1 |
| Q192R | 16,91<br>(13,98 – 20,47) | ~1,9 |
| Q192R+F305L | 27,80<br>(21,67 – 35,69) | ~3,1 |

## Acknowledgements

E.H. is a recipient of a DOC Fellowship of the Austrian Academy of Science. We thank Univ. Prof. Dr. Florian Krammer, Anna Fürst, MMSc., Ivan Ploner, MSc. and Seyad Arad Mogadashi, MSc. for useful discussions and technical assistance.

## Author contributions

E.H. conceived the initial concept. E.H., A.V. and F.C. designed the experiments. E.H. conceived the cloning strategies. E.H. generated recombinant viruses. A.V. previously established a reliable lentiviral transduction system in our institute. E.H., A.V. and F.C. performed experiments. E.H. and D.v.L. wrote the manuscript. D.v.L. and E.H. provided the infrastructure and funding to the project. D.v.L. provided supervision of E.H. All authors read and approved the final manuscript.

## Data availability

All pertinent data to support this study are included in the manuscript and supplementary files. If required, further data supporting the findings are available upon request.

